# Phylogenetic analysis of SARS-CoV-2 lineage development across the first and second waves in Eastern Germany, 2020

**DOI:** 10.1101/2021.04.29.441906

**Authors:** Buqing Yi, Anna R. Poetsch, Marlena Stadtmüller, Fabian Rost, Sylke Winkler, Alexander H. Dalpke

**Affiliations:** Institute of Medical Microbiology and Virology, Medical Faculty, Technische Universität Dresden, Dresden, Saxony, Germany; Biotechnology Center (BIOTEC), Technische Universität Dresden, Dresden, Saxony, Germany; National Center for Tumor diseases (NCT), Dresden, Saxony, Germany; DRESDEN concept Genome Center, Technische Universität Dresden, Dresden, Saxony, Germany; Center for Regenerative Therapies Dresden, Technische Universität Dresden, Dresden, Saxony, Germany; Max Planck Institute of Molecular Cell Biology and Genetics, Dresden, Germany and DRESDEN concept Genome Center, Technische Universität Dresden, Dresden, Saxony, Germany

**Keywords:** SARS-CoV-2, COVID-19, phylogenetic analysis, second wave

## Abstract

SARS-CoV-2 lineages prevalent in the first and second waves in Eastern Germany were different, with many new variants including four predominant lineages in the second wave, having been introduced into Eastern Germany between August to October 2020, indicating the major cause of the second wave was the introduction of new variants.

---

In Germany, the first wave of the coronavirus disease (COVID-19) pandemic (March to May, 2020) showed visible regional differences: it was much milder in Eastern regions (Saxony, Saxony-Anhalt, Berlin, Brandenburg and Thuringia) compared to most other regions in Germany. However, the severity of the second wave (August to December, 2020) was similar in most regions in Germany. It is unclear how the second wave started in Eastern Germany where in June and July 2020 the number of COVID-19 cases was close to zero (Fig. 1A). We therefore performed phylogenetic analysis of the predominant SARS-CoV-2 variants in the first and second waves in Eastern Germany. By dissecting the difference between the first wave and the second wave, we expect the information achieved through this study could provide insights into the cause of the second wave and can possibly help developing suitable strategies for preventing similar scenarios in future.

**Fig. 1.**
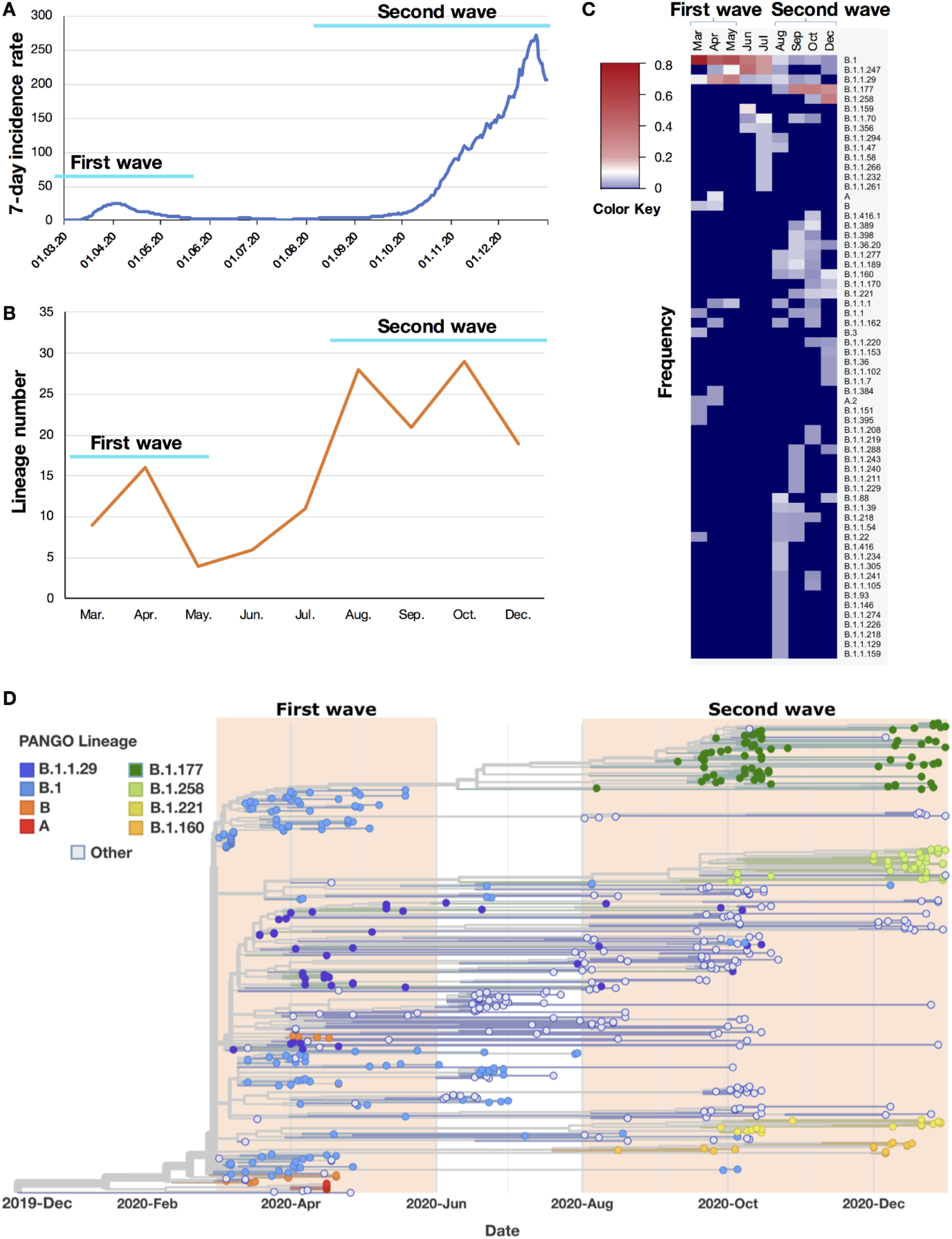
Analysis of SARS-CoV-2 lineages predominant in the first and second wave in Eastern Germany, March to December 2020. **A**. 7-day incidence rate per 100,000 inhabitants in Eastern Germany. First wave: March to May; Second wave: August to December. **B**. Summary of detected total SARS-CoV-2 lineage numbers in each month. C. Frequency of detection for each SARS-CoV-2 lineage in each month in Eastern Germany (range: 0 ∼ 0.74, representing 0% ∼ 74%; 0 is shown with deep blue, indicating no detection of the relevant variant in that month). Note: to achieve a better resolution, a few lineages that were detected only once across 2020 and with a frequency < 0.01 were omitted from the heatmap. D. Phylogenetic and time tree of SARS-CoV-2 genomes from Eastern Germany, March to December 2020. Each genome is denoted with Pangolin-lineage (PANGO Lineage). The names of lineages that were predominant in the first or second wave are colour labelled. The four lineages from the second wave B.1.177, B.1.258, B. 1.221 and B.1.160 had been circulating in multiple other European countries since June (5).

For surveillance purpose, randomly selected SARS-CoV-2 positive samples from each state in Germany were sequenced by the Robert Koch Institute or by sequencing facilities of local universities. All sequences that passed stringent quality control were uploaded to GISAID (1). We used GISAID sequences from regions in Eastern Germany dating between March to December 2020 in this study (data collected on Feb. 28, 2021; Table S1). The number of genomes in each month was: 74 (Mar), 102 (Apr), 19 (May), 48 (Jun), 18 (Jul), 41 (Aug), 47 (Sep), 105 (Oct), 112 (Dec) (only a few genomes were sequenced in November because the testing labs were extremely overloaded by then, so the data of November was not included in the analysis). The data of 7-day-incidence rate per 100,000 inhabitants was obtained for the states in Eastern Germany from the Robert Koch Institute (https://www.rki.de/DE/Content/InfAZ/N/Neuartiges_Coronavirus/Daten/Fallzahlen_Daten.htm), and the average values were visualized in Fig. 1A. Lineage group assignment of SARS-CoV-2 genomes was performed with software package **P**hylogenetic **A**ssignment of **N**amed **G**lobal **O**utbreak **LIN**eages (Pangolin) (2). Phylogenetic maximum likelihood (ML) and time trees were constructed using the SARS-CoV-2-specific procedures taken from github.com/nextstrain/ncov (3, 4).

The first wave in Eastern Germany reached its peak in April 2020 (Fig. 1A). Based on the frequency of detection in April (Fig. 1C&D), the SARS-CoV-2 lineages predominant in the first wave were: B.1, B.1.1.29, A and B, with respective frequencies of 46%, 21%, 9% and 7% (shown as 0.46, 0.21, 0.09 and 0.07 in Fig. 1C). The second wave reached its peak in December 2020 (Fig. 1A). Based on the frequency of detection in December (Fig. 1C&D), the most prevalent lineages in the second wave were different from that of the first wave: B.1.258, B.1.177, B.1.160 and B.1.221, with respective frequencies of 32%, 25%, 9% and 7%. All lineages in the first and second waves were defined in one batch with the pangoLEARN_version 2021-02-21. These four lineages B.1.258, B.1.177, B.1.160 and B.1.221 from the second wave were neither detected in the first wave in Eastern Germany (Fig. 1C&D), nor possibly derived from the local first wave lineages through mutant accumulation since the 7-day incidence rate in June and July in Eastern Germany was close to zero, which means there was almost no virus circulating in the local population. B.1.258, B.1.177, B.1.160 and B.1.221 were first identified in other European countries before April 2020 (https://cov-lineages.org/pango_lineages.html), and have a known spreading history in multiple other European countries in June and July, such as in Spain (5). In Eastern Germany B.1.258 was first detected in October; B.1.177 was first detected in August; B.1.160 was first detected in August; B.1.221 was first detected in September (Fig. 1C&D).

From August till October 2020 was the summer/autumn holiday season in Eastern Germany, and a lot of regional and international travelling took place during this period. Our analysis indicates that many new lineages were introduced into Eastern Germany from August to October 2020 (Fig. 1B&C). For example, in August, 20 new lineages were first detected in Eastern Germany, such as B.1.160, B.1.1.234, B.1.1.277, B.1.1.305, B.1.1.39, B.1.416 and B.1.177. In total, more than 40 new variants were introduced into Eastern Germany during the holiday season (Fig. 1C), including the four predominant lineages B.1.258, B.1.177, B.1.160 and B.1.221, which paved the base for the second wave.

Interestingly, only a few of these new variants were responsible for most local cases in December when the second wave reached its peak value: the four predominant new variants (B.1.258; B.1.177; B.1.160; B.1.221) were estimated to account for more than 70% of the cases based on their frequency of detection. These findings suggest that many control measures, such as test on the airport, might have prevented the local transmission of many new variants. However, in August to October 2020, the lineages B.1.258, B.1.177, B.1.160 and B.1.221 were prevalent in many European countries (5), which means the chances of the introduction of these lineages were higher than other variants. In addition, travelling within Germany might also play a role here since no tests were required for regional travelers.

In conclusion, the introduction of various SARS-CoV-2 lineages from August to October 2020 was the major driving force for the development of the second wave in Eastern Germany regions, instead of expansion of local circulating lineages from the first wave.

## Acknowledgments

We thank all researchers who are working around the clock to generate and share genome data on GISAID (http://www.gisaid.org) on which the analysis is based. We specifically thank colleagues at the Institute of Medical Microbiology and Virology, Technische Universität Dresden, for their work in performing SARS-CoV-2 sample testing and sequencing sample preparing, and we thank the Robert Koch Institute and Dresden Concept Genome Center for their sequencing efforts.

## Declarations

No potential conflict of interest was reported by the author(s).

## Availability of data

The data used in this work is publicly available (1).

## Notes

### Competing Interest Statement

The authors have declared no competing interest.

